# Phylogenomics unravels the early diversification of fungi

**DOI:** 10.1101/2021.12.12.472261

**Authors:** Jürgen F. H. Strassert, Michael T. Monaghan

**Author notes:** Correspondence (J.F.H.S.).

## Abstract

Phylogenomic analyses have boosted our understanding of the evolutionary trajectories of all living forms by providing continuous improvements to the tree of life^1–5^. Within this tree, fungi represent an ancient eukaryote group^6^, having diverged from the animals ∼1.35 billion years ago^7^. Estimates of the number of extant species range between 1.5 and 3.8 million^8,9^. Recent reclassifications and the discovery of the deep-branching Sanchytriomycota lineage^10^ have brought the number of proposed phyla to 20^11^; 21 if the Microsporidia are included^12–14^. Uncovering how the diverse and globally distributed fungi are related to each other is fundamental for understanding how their lifestyles, morphologies, and metabolic capacities evolved. To date, many of the proposed relationships among the phyla remain controversial and no phylogenomic study has examined the entire fungal tree using a taxonomically comprehensive data set and suitable models of evolution. We assembled and curated a 299-protein data set with a taxon sampling broad enough to encompass all recognised fungal diversity with available data, but selective enough to run computationally intensive analyses using best-fitting models. Using a range of reconstruction methods, we were able to resolve many contested nodes, such as a sister-relationship of Chytridiomyceta to all other non-Opisthosporidia fungi (with Chytridiomycota being sister to Monoblepharomycota + Neocallimastigomycota), a branching of Blastocladiomycota + Sanchytriomycota after the Chytridiomyceta but before other non-Opisthosporidia fungi, and a branching of Glomeromycota as sister to the Dikarya. Our most up-to-date fungal tree of life will serve as a springboard for future investigations on the evolution of fungi.

## Results

### A well-curated and taxon-balanced phylogenomic data set

To untangle the early diversification in the fungal tree of life, we assembled a 299-protein data set with 661 taxa that represent all extant recognised fungal lineages (Table S1). All proteins were carefully inspected by reconstructing single-protein phylogenies to evaluate orthology and detect contamination. Considering that the taxonomic distribution of publicly available fungal genomes and transcriptomes is highly biased towards Ascomycota and Basidiomycota, we placed emphasis on the inclusion of underrepresented phyla such as the Blastocladiomycota or the fast-evolving early-branching Aphelidiomycota, Rozellomycota, and Microsporidia (together known as Opisthosporidia). To uncover deep phylogenetic relationships, a taxon-reduced data set (101 taxa) was derived to allow for more computationally intensive analyses. This taxon-reduced data set maintained a high diversity and coverage and selected slow-evolving over fast-evolving lineages within phyla.

### Reconstructing the phylogeny of fungi

Two trees were inferred from the full data set, one by maximum likelihood (ML) under the site-homogeneous LG+G+F model, and the other under the multi-species coalescent (MSC) model. These were largely congruent to each other (Figures S1 and S2); however, as expected for the heterogeneous taxon sampling in our full data set, some of the deeper nodes remained unsupported using these models. The taxon-reduced data set, from which all final analyses were derived and to which is referred to hereafter, was therefore analysed under two site-heterogeneous mixture models: CAT+GTR+Γ4 using Bayesian inference (BI) (hereafter *CATGTR*; Figure 1A), and LG+C60+G+F with posterior mean site frequency profiles using ML (hereafter *ML-C60*; Figures 1B and S3). Both models have been shown to better account for rate variation across sites^15,16^ and their implementation led to the recovery of a similar overall topology with maximal statistical support for nearly all nodes (Figure 1A, B). A further tree was inferred under the MSC model, which was highly congruent to the *ML-C60* tree (Figures 1B, S3, and S4). All three analyses (*CATGTR, ML-C60*, and MSC; Figure 1, Table 1) were congruent regarding the branching of Chytridiomyceta as sister to all other non-Opisthosporidia fungi; the sister-relationship of Chytridiomycota to a clade comprising Neocallimastigomycota + Monoblepharomycota; the branching of Blastocladiomycota + Sanchytriomycota, which diverged after the Chytridiomyceta but before other non-Opisthosporidia fungi; the sister-relationship of *Olpidium* to all other non-zoosporic fungi; or the sister-relationship of Pucciniomycotina to Agaricomycotina/Wallemiomycotina + Ustilaginomycotina. Aphelidiomycota was fully supported as sister taxon to the entire non-Opisthosporidia clade and distinct from Rozellomycota and Microsporidia, although Aphelidiomycota was represented by only one species in our analysis (Figure 1).

**Table 1.**
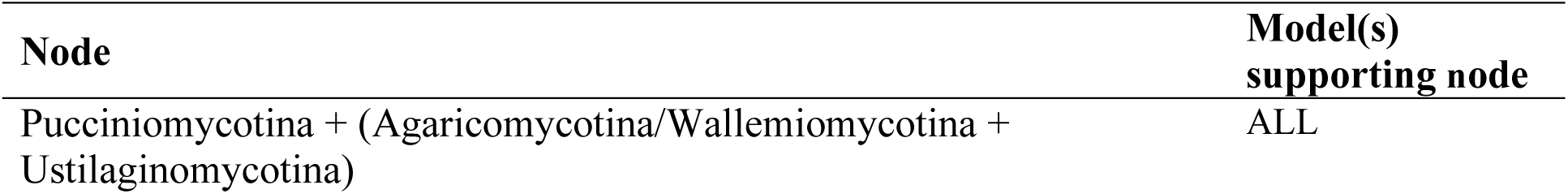
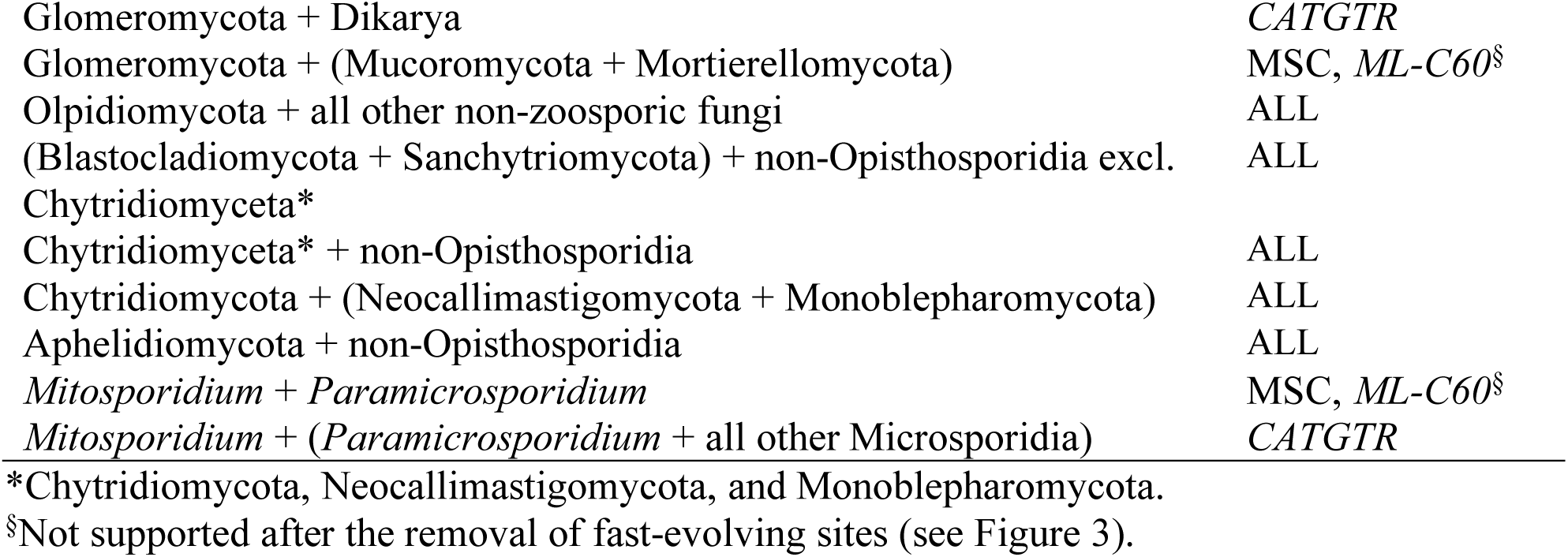
Nodes of major interest.

**Figure 1.**
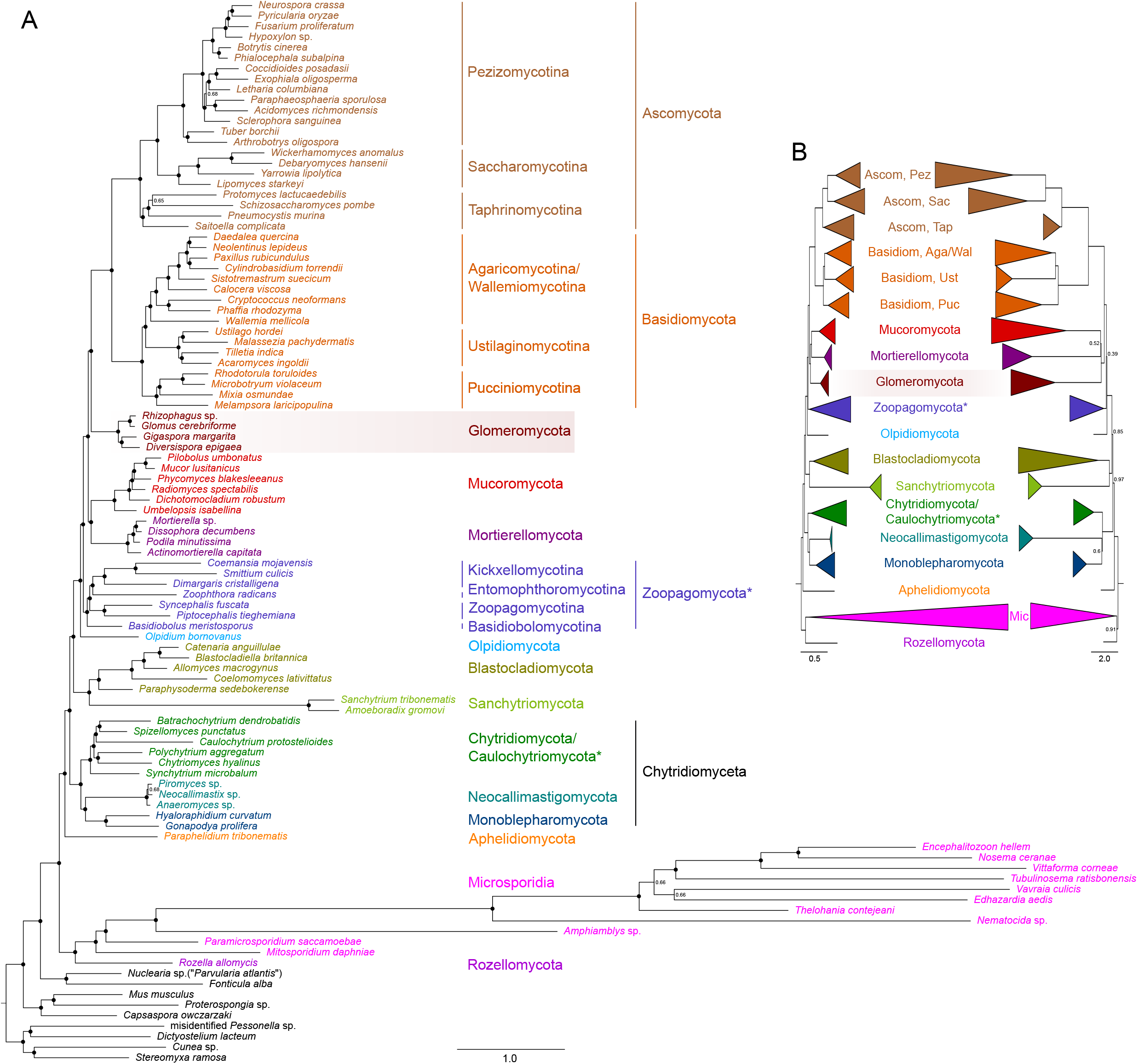
Phylogenetic trees showing the early diversification of fungi. (A) Bayesian consensus tree inferred from a concatenated alignment of 299 proteins and 101 taxa under the CAT+GTR+Γ4 (*CATGTR* in the text) model. Node support is given by posterior probabilities (full support is indicated by black circles). (B) Two phylogenetic trees showing perfect congruence. Left: Tree inferred by ML under the LG+C60+G+F-PMSF (*ML-C60* in the text) model using the same alignment as used for the Bayesian tree. Nonparametric bootstrap support is not indicated but was maximal for all nodes shown (for details, see Figure S3). Right: Tree based on summary-coalescent analyses of 299 single-protein ML phylogenies. Quadripartition support values are given for nodes that are not fully supported (for details, see Figure S4). The conflicting branching of the Glomeromycota in the CAT+GTR+Γ4 analysis and the LG+C60+G+F-PMSF and summary-coalescent analyses is highlighted. \**Caulochytrium* and the subphyla of the Zoopagomycota were recently proposed to be independent phyla^11^ (see Discussion).

Ambiguity remained considering the diverging pattern of the Glomeromycota, either as sister to Mucoromycota + Mortierellomycota as reconstructed in the MSC and *ML-C60* trees or as sister to the Dikarya (Ascomycota + Basidiomycota, and possibly Entorrhizomycota that lack genomic/transcriptomic data^12^) as reconstructed in the *CATGTR* tree (Figure 1). Another discrepancy between the inferred tree topologies was the placement of the opisthosporid *Mitosporidium* as either sister to only *Paramicrosporidium* (MSC and *ML-C60*; Figures S3 and S4) or sister to all other Microsporidia including *Paramicrosporidium* (*CATGTR*; Figure 1A).

### Tree evaluation

The conflicts in the branching of the Glomeromycota and *Mitosporidium* were fully supported by bootstrap (*ML-C60*) and posterior probability (*CATGTR*) analyses. Both alternative branching orders in the *CATGTR* topology were also rejected by ML in Approximately Unbiased tests (Glomeromycota + Dikarya, p-AU = 0.022; *Mitosporidium* + all other Microsporidia, p-AU = 0.011). To test whether the conflicts were due to the usage of different tree reconstruction methods (ML or BI), a further tree was derived using BI under the same C60 model that was used for the ML tree. This tree (hereafter *BI-C60*; Figure S5) recapitulated the topology obtained with *ML-C60* indicating that differences were due to the evolutionary model rather than the inference method. To test which model better minimised inadequacy in describing compositional heterogeneity, posterior predictive analyses were conducted that resulted in a better fit of *CATGTR* compared to *BI-C60* (Table 2). We also removed 25% and 50% of the most compositionally heterogeneous sites from the alignment in order to reduce compositional heterogeneity, and additional trees were reconstructed from these new alignments with *CATGTR* (Figure 2). The inferred trees were in agreement with the *CATGTR* tree reconstructed from the full-length alignment (Figures 1A and 2).

**Table 2.** Results of the posterior predictive analyses (PPA).

| Model | Site-specific AA heterogeneity* |  |  | Lineage-specific AA heterogeneity* |  |
| --- | --- | --- | --- | --- | --- |
|  | div | conv | var | max | mean |
| CAT+GTR+G | 7.93110 | 13.16270 | 13.58923 | 179.34 | 340.850 |
| LG+C60+G | 75.71623 | 46.49947 | 44.97260 | 270.14 | 472.514 |

\*z-scores

**Figure 2.**
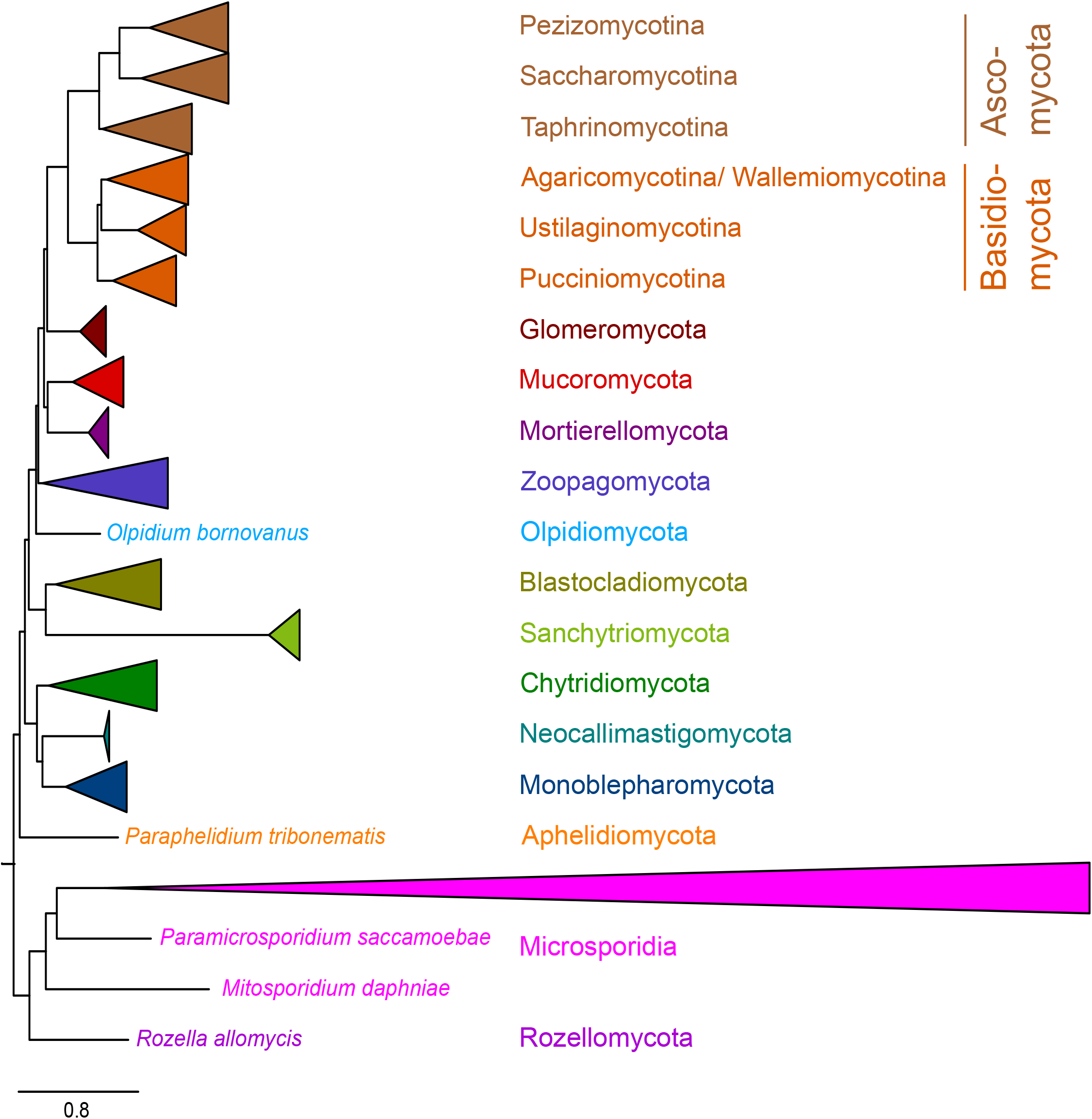
Heterogeneous sites removal analysis. Simplified Bayesian consensus tree inferred under the CAT+GTR+Γ4 (*CATGTR* in the text) model from a concatenated alignment of 299 proteins and 101 taxa with 50% of the most heterogeneous sites removed. All nodes shown were fully supported. A further tree, which was inferred from the same alignment but with 25% of the most heterogeneous sites removed, had an identical topology (not shown). Microsporidians are only partly collapsed in order to depict the position of *Paramicrosporidium saccamoebae. Caulochytrium protostelioides* was nested within the Chytridiomycota (same branching pattern as in Figure 1A; see Discussion).

In order to understand the extent to which fast-evolving sites, which are more likely to be homoplasic, led to systematic bias in the *ML-C60* tree inference due to model misspecification^17^, the fastest-evolving sites were progressively removed from the alignment in increments of 10,000 sites. Each sub-alignment was then subjected to further ML tree inferences using the LG+C60+G+F model and node support was inferred by ultrafast bootstrap approximation. Because of the higher computational demand of BI, this analysis was run for ML inferences only. Interestingly, whereas the bootstrap support for Dikarya remained maximal throughout all the analyses, the support for the sister-relationship of Glomeromycota to Mucoromycota + Mortierellomycota decreased rapidly while the support for their closer relationship to the Dikarya, as seen in the *CATGTR* trees reconstructed from complete and pruned alignments, increased, reaching 92% after the removal of 30,000 sites (Figure 3). Further removal of sites led either to an unsupported branching or recovered the Glomeromycota + Dikarya topology again (100,000 sites removed; Figure 3). For the fast-evolving microsporidians, the sister-relationship of *Mitosporidium* to all other Microsporidia, as inferred by *CATGTR*, was only supported when the 120,000 fastest-evolving sites were removed from the alignment (i.e. 8,450 sites remained; Figure 3).

**Figure 3.**
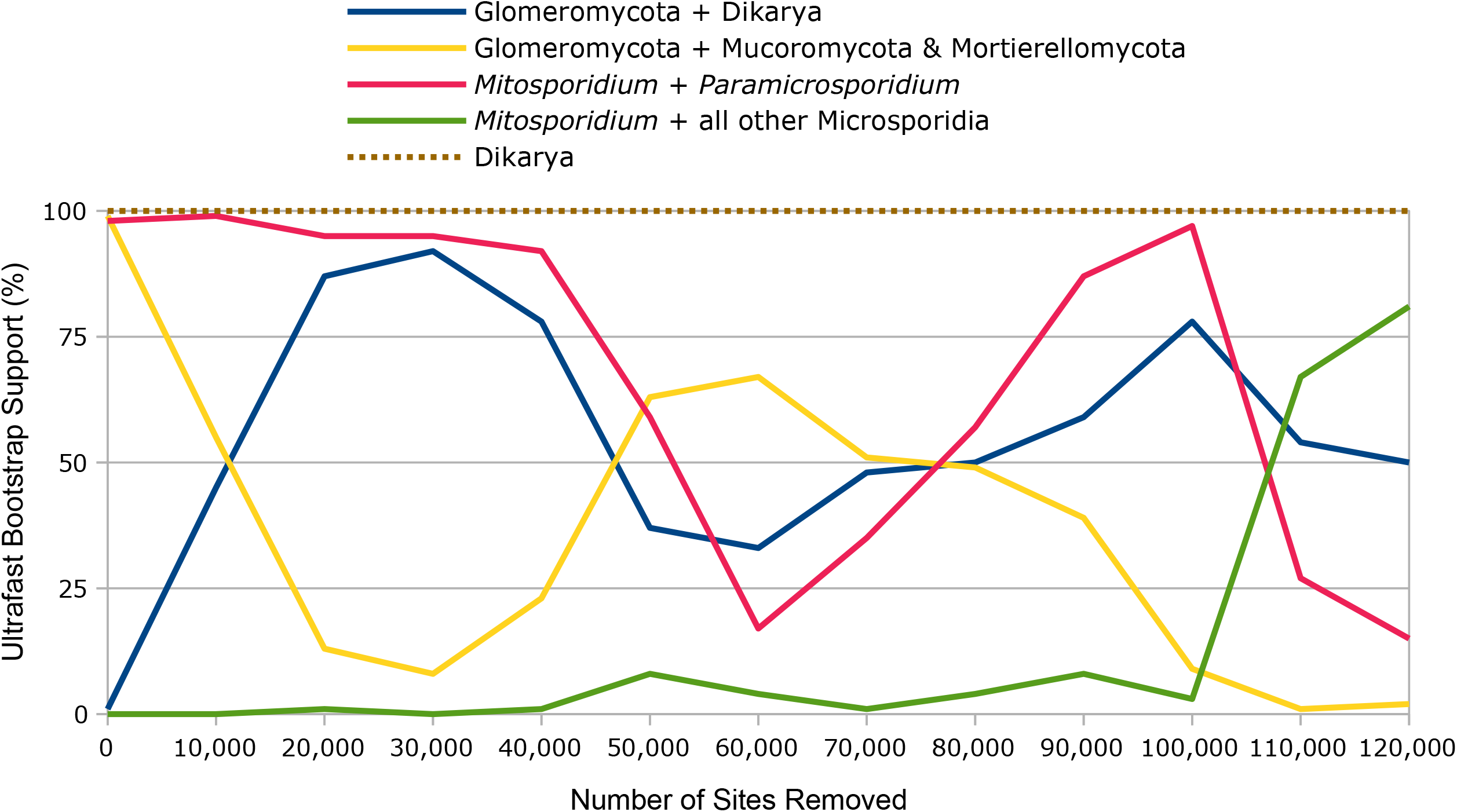
Fast-evolving sites removal analysis. ML ultrafast bootstrap approximation for selected nodes using the full matrix and subsets from which fast-evolving sites were removed in 10,000 increments.

Finally, to assess whether the here received topologies were skewed due to long branch attraction (LBA) effects^18^, further *ML-C60* and *CATGTR* trees were calculated but with the Sanchytriomycota and the nine fastest-evolving Microsporidia species removed. In both trees, the branching order of the remaining fungal groups was fully congruent to the branching order that has been recovered before under the same evolutionary model, using the alignment that includes the fast-evolving taxa (compare Figure 1 with Figure S6).

## Discussion

Fungi show a great diversity in morphology and lifestyle. They are important decomposers and through the provision of nutrients, various lineages are essential for the survival of >80% of all plant species^19^. As parasites, chytrid infections can have a significant impact on amphibian populations^21^ and may even affect the planet’s carbon cycle by terminating algal blooms^20^. Despite their ecological importance, worldwide distribution, and essential roles in human history as food source, leavening and fermentation agents, and medicines, our knowledge of how the huge diversity of fungi evolved is still limited. Phylogenomic studies encompassing the entire fungal kingdom with nearly all its different phyla are scarce — either due to missing data or to a focus on only a few certain groups of fungi^10,22–25^. In this study, we tested whether a more balanced taxon-sampling, in combination with new filtering methods^26,27^ (see STAR*Methods) that improve the phylogenetic signal of each single-protein alignment and best-fitting models of evolution, could untangle some of the hitherto debated early diversifications in the fungal tree of life.

The branching order among fungal phyla remained consistent and well supported across multiple analyses using site-heterogeneous models in BI (*CATGTR* and *BI-C60*) and ML (*ML-C60*) tree reconstructions, and using an MSC approach (Figures 1, 2, and S3–S6). This applies even to phyla that were represented by only a single species such as Olpidiomycota, whose sister-relationship to non-flagellated fungi has recently been proposed based on phylogenomics^28^. The sole exception (on the phylum level) concerned the placement of the Glomeromycota, which was sister to the Dikarya in *CATGTR*-based analyses or was sister to the Mucoromycota/Mortierellomycota clade in the other analyses. Glomeromycota were suggested to form an independent phylum at the beginning of this century^29^, but closer phylogenetic affiliation has remained controversial ever since^12,25,30,31^. A sister-relationship to the Dikarya has been proposed based on tree inferences using the elongation factor EF-1 alpha, RNA polymerase II subunits and/or rRNA gene alignments^6,29,31–33^. This placement was discussed to be artificial due to lineage-specific rates in these data sets^12^ and was rejected in a ML tree inference from a 192-protein matrix. However, we note that this matrix analysis used an LG model with fixed base frequencies^34^, and that our results support a more recent *CATGTR*-based analysis of a 264-protein matrix that also recovered Glomeromycota as sister to the Dikarya^10^. It is reasonable to assume that this finding depicts the real scenario in the evolution of fungi and that the discovered sister-relationship to the Mucoromycota/Mortierellomycota clade in the *ML-C60* tree is a consequence of the inadequacy of the C60 model for capturing compositional heterogeneity. This is in line with the results obtained by the posterior predictive test, in which the fit of both models, *CATGTR* and *BI-C60*, was directly compared using the PhyloBayes software package (Table 2). Congruent topologies reconstructed from the analyses of the same alignment using both *BI-C60* and *ML-C60* (Figures S3 and S5) give further evidence that the found discrepancies in the branching order of the Glomeromycota are rather caused by the less fitting evolutionary C60 model than by the use of BI or ML. Interestingly, however, Glomeromycota + Dikarya was also supported by *ML-C60* tree inferences after only 15% of the fastest-evolving sites were removed (Figure 3), indicating that the conflicting placement of Glomeromycota in the C60 analysis of the full alignment may be due to statistical problems this model encounters with homoplasic sites.

The relationships among the opisthosporidian phyla remained controversial for a long time^6,22,35,36^. Here, the Aphelidiomycota representative was placed as sister to all non-Opisthosporidia fungi in all analyses (Figures 1, 2, and S3–S6). Similar findings have been reported before based on phylogenomic studies corroborating the paraphyletic nature of Opisthosporidia^10,35^. The assignment of some of the earliest-branching fungal species to either the Rozellomycota or the Microsporidia is more controversial, leading to debates as to whether these phyla are monophyletic or not^6,11,22^. *Paramicrosporidium saccamoebae* was initially assigned to the Rozellomycota based on a phylogenetic analysis of SSU rRNA gene sequences, but an unsupported affiliation to the Microsporidia was recovered when 5.8S and partial LSU rRNA gene sequences were added to the analyses^37^. We found the exact placement of *P. saccamoebae* to be questionable. While trees inferred with the better fitting *CATGTR* model recovered this species to be nested between the microsporidians *Mitosporidium daphniae* and *Amphiamblys* sp. (Figures 1A and 2), a sister-relationship to *M. daphniae* was recovered in the MSC tree and with the C60 model in BI and ML inferences (Figures S3–S5). For these fast-evolving lineages, the latter topology was only challenged when at least 85% of the fastest-evolving sites were removed (Figure 3). However, independent from the exact position of *P. saccamoebae*, the only safely assigned Rozellomycota species with available genomic data, *Rozella allomycis*, formed a sister to all Microsporidia and *P. saccamoebae* in all analyses (Figures 1, 2, and S3–S6). Taking into account these results, the results obtained by rRNA analyses^37^ (see above) and the morphology of *P. saccamoebae*, which resembles that of microsporidians (they lack a mitochondrion and have tiny spores with a polar filament^37^), its assignment to the Microsporida seems to be reasonable and is proposed here. Thus, Microsporidia and Rozellomycota form two distinct lineages for which a frequently discussed paraphyly^6,22,25^ cannot be confirmed here. Whether or not Microsporidia + Rozellomycota should be classified as only one phylum is currently debated^6,31,38^ and will require data from additional representatives of Rozellomycota.

Throughout all analyses, our study revealed not only consistency regarding the previously contested phylogenetic position of the zoosporic subkingdom Chytridiomyceta^e.g. 10,25^, which live either as saprophytes, parasites, or as intermediate forms, but also regarding the discussed branching order among its phyla^6,12,30^. With that, however, the justification for the separation of the chytridiomycete phylum “Caulochytriomycota” from the Chytridiomycota remains enigmatic (the erection of this phylum was proposed by Doweld in 2014^39^ and has been adapted by other researchers since). Only two species of the genus *Caulochytrium* are known, and DNA sequences are available only for *Caulochytrium protostelioides*. In our analysis, *C. protostelioides* was either closely related to, or even nested within, the Chytridiomycota, and the clade was well-supported in all analyses (Figures 1, 2, and S3–S6). We therefore suggest *Caulochytrium* be assigned to the Chytridiomycota based on current data. The here unambiguously documented sister-relationship of Chytridiomycota to Neocallimastigomycota + Monoblepharomycota makes them the earliest diverging group within the Chytridiomyceta — a finding that is rather surprising taking into account that the zoospores of Chytridiomycota show more morphological similarities to those of Monoblepharomycota than to those of Neocallimastigomycota (in addition, members of the latter are the only opisthokonts, whose zoospores can bear multiple flagella)^6,12^. Our study is the first to position the chytrid *Hyaloraphidium curvatum* based on phylogenomic analyses and clearly shows its affiliation to the Monoblepharomycota, which was represented by *Gonapodya* in our data set. *Hyaloraphidium* was initially miss-classified as a green alga and later proposed to cluster with Chytridiomyceta based on SSU rRNA gene sequence analysis^40^. However, its closer affiliation has remained enigmatic ever since.

The previously unclear^12,25^ but here consistent placement of Blastocladiomycota, which together with the Sanchytriomycota diverged after the Chytridiomyceta but before other non-Opisthosporidia fungi, seems in agreement with the finding of a distinct morphology of their zoospores from those produced by Chytridiomyceta. Another distinguishing feature is the sporic meiosis that results in a regular alternation of haploid and diploid generations in Blastocladiomycota^41^.

### Final remarks

The here presented tree (Figure 1A) provides a robust backbone topology for the diversification of major fungal groups. For the recovery of ancient nodes, not only site rate variation but also variation of the sequence composition across branches and/or alignment sites can hamper the reconstruction of topology and branch length. Thus, 25% and 50% of the sites that contribute the most to branch heterogeneity were removed from the alignment and new *CATGTR* trees were inferred (Figure 2). Both trees reflected the overall topology of the tree inferred from the complete alignment (Figure 1A) indicating that compositional biases had either no or a negligible negative impact that could result in model misspecification despite using the best-fitting CAT+GTR+Γ4 model. Considering that also LBA effects did not seem to have a significant impact on the received *ML-C60* and *CATGTR* topologies (Figures 1 and S6), it is likely that the found incongruence between these trees resulted from an inability of the *ML-C60* model to model fast-evolving and heterogeneous sites. This would be in line with the findings of the fast-evolving sites removal analysis (Figure 3). However, it is noteworthy that the *CATGTR* backbone topology provided here — like all phylogenetic trees — represents a hypothesis and it is safe to assume that with an increasing diversity of fungi for which genomic/transcriptomic data will be available, several clades will be reshuffled. It should also be noted that the taxonomic classification in fungi is changing rapidly. The lineage information in this study was assigned according to the NCBI taxonomy (ncbi.nlm.nih.gov/taxonomy) with some modifications especially regarding the phylum assignments, which, if not otherwise noted, were adopted in order to match those recently proposed by Wijayawardene *et al*.^11^ A comprehensive revision of the taxonomic classification in fungi, based on this and other recent studies^e.g. 10,25^, is desirable.

## Supplemental information

Table S1: Lineage and source information for all taxa used in this study.

Document S1: Figures S1–S6.

Data S1: Sequence data, single-protein trees, and BI consensus trees from individual MCMC chains.

## Acknowledgments

J.F.H.S. acknowledges support from the German Research Foundation (DFG; Grant STR1349/2-1 Project No. 432453260) and research was partially funded by the German Federal Ministry of Education and Research (BMBF, Förderkennzeichen 033W034A). The authors thank the High-Performance Computing Service of ZEDAT, Freie Universität Berlin, for computing time. Finally, we thank David Moreira and Luis Javier Galindo for sharing sequence data for *Amoeboradix gromovi* and *Sanchytrium tribonematis*.

## Author contributions

J.F.H.S. conceived the study, assembled and curated the data, performed all analyses, and drafted the manuscript. M.T.M. contributed to the final manuscript. Both authors read and approved the final version.

## STAR*Methods

### Key resources table

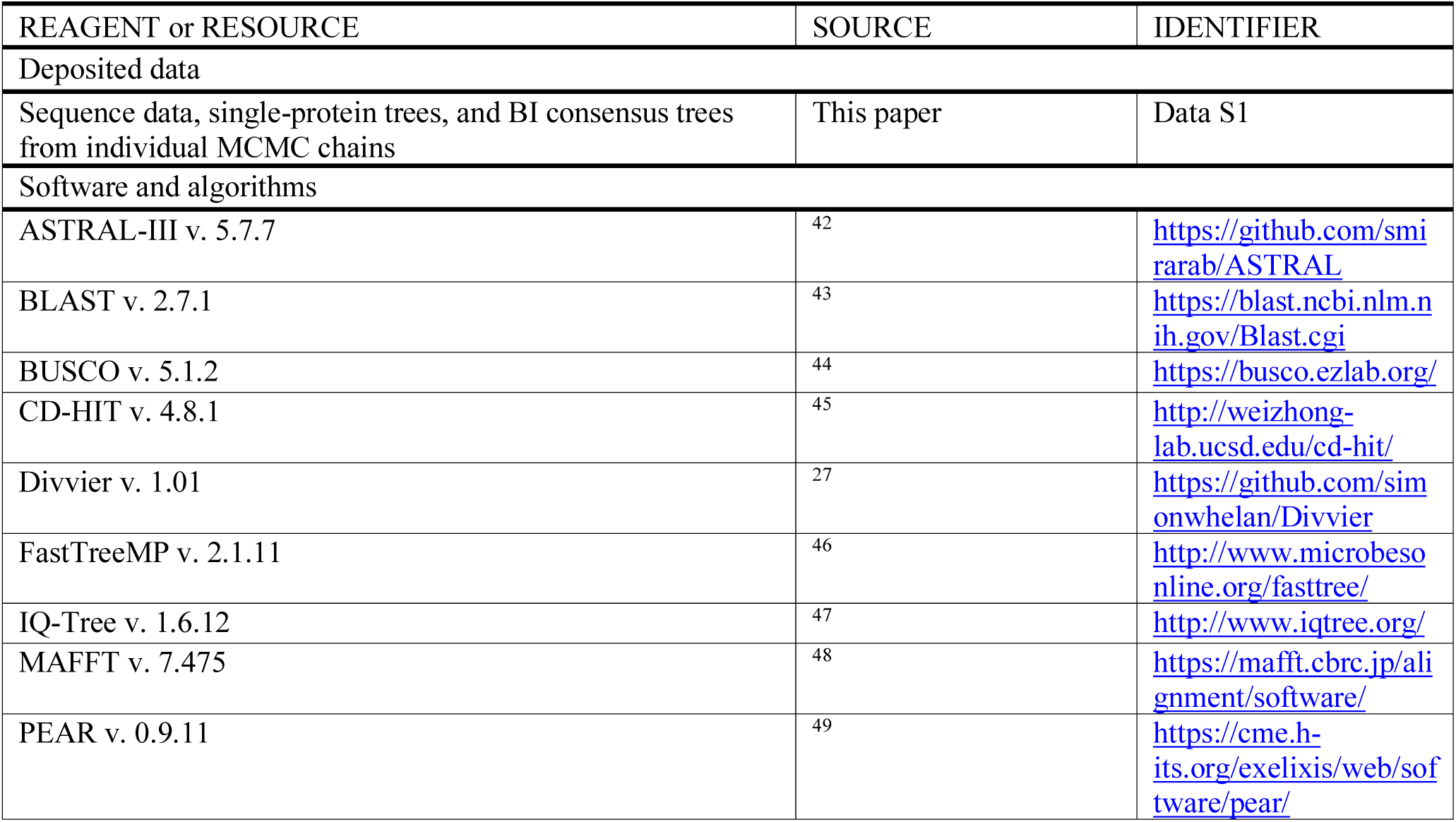
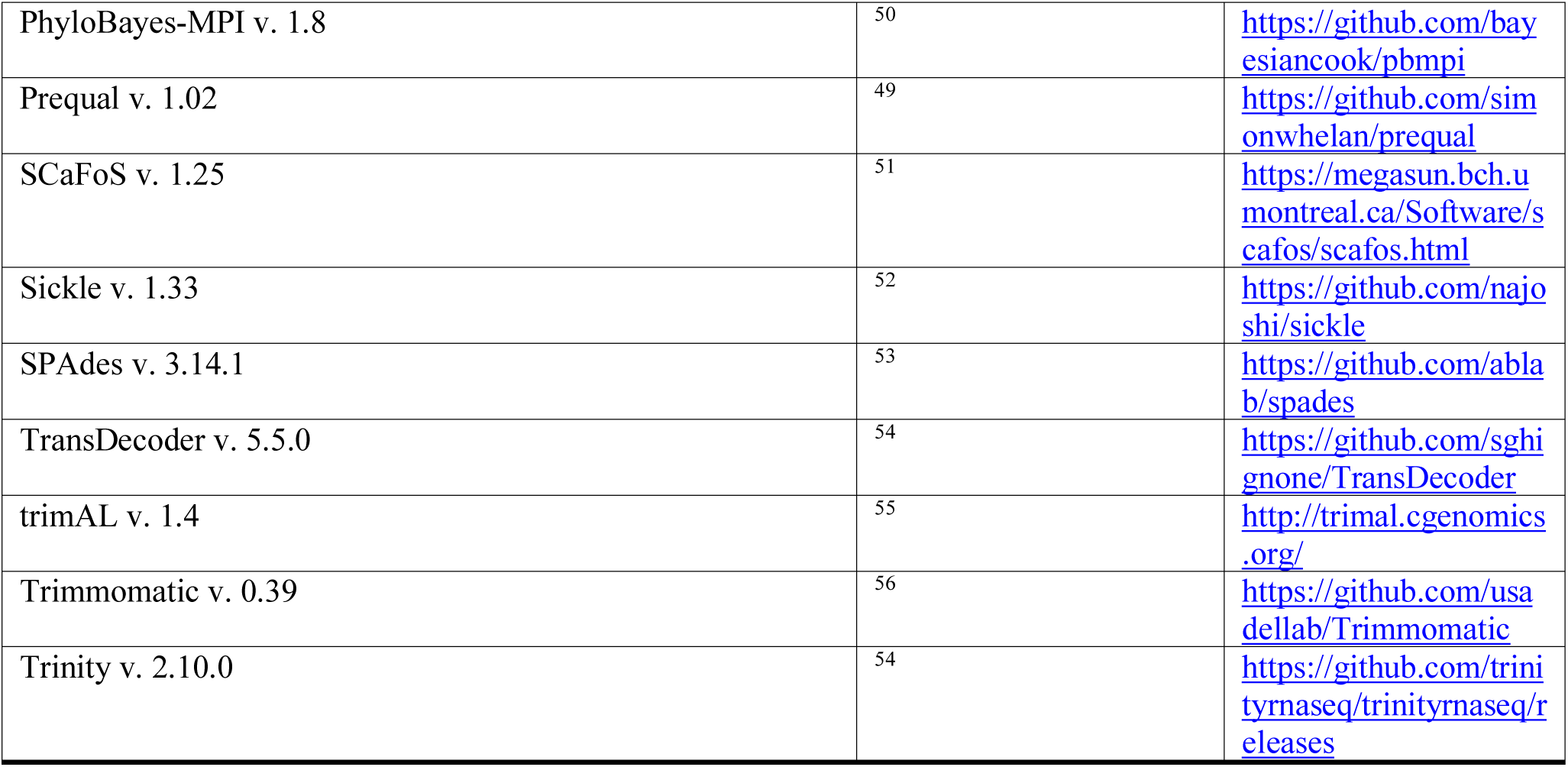

### Resource availability

#### Lead contact

Further information and requests for resources should be directed to and will be fulfilled by the lead contact, Jürgen F. H. Strassert.

### Materials availability

This study did not generate new unique reagents.

### Data and code availability

All data needed to evaluate the conclusions of this study are presented in the paper and the Supplemental information. Raw sequence data are available under the following web-links: doi.org/10.6084/m9.figshare.12417881.v2, fungi.ensembl.org, metazoa.ensembl.org, protists.ensembl.org, imicrobe.us/#/projects/104, mycocosm.jgi.doe.gov, and ncbi.nlm.nih.gov. The script alignment_pruner.pl is available at github.com/novigit/davinciCode/blob/master/perl. Any additional information required to reanalyse the data reported in this paper is available from the lead contact upon request.

### Experimental model and subject details

Genomes and transcriptomes were downloaded from EukProt, Ensembl, Imicrobe, Mycocosm, and NCBI. Sequences of outgroup taxa were added from Strassert et al.^7^ For detailed source information, see Table S1. Taxa were selected such that all major fungal groups were represented, neglecting taxa with more missing data. Analyses were carried out using the Zedat high-performance computing infrastructure at Freie Universität Berlin^57^.

### Method details

#### Phylogenomic data set construction

Genomes and transcriptomes of a broad range of fungi and selected non-fungal lineages were downloaded from EukProt^58^, fungi.ensembl.org, metazoa.ensembl.org, protists.ensembl.org, imicrobe. us/#/projects/104, mycocosm.jgi.doe.gov, and ncbi.nlm.nih.gov (Table S1). For unassembled sequence data, paired end reads were trimmed using Trimmomatic v. 0.39^56^ with the following flags employed: LEADING: 3, TRAILING: 3, SLIDINGWINDOW: 4:15, and MINLEN: 36. Transcriptomic data was assembled and translated into amino acids by using the default settings in Trinity v. 2.10.0 and TransDecoder v. 5.5.0, respectively^54^. Reads of genomic data were merged with PEAR v. 0.9.11^49^, quality filtered with Sickle v. 1.33^52^ (only the paired but unmerged reads), and assembled with SPAdes v. 3.14.1^53^ using the --isolate option. Genes were predicted using BUSCO v. 5.1.2 with the database fungi_odb10^44^.

A recently published data set comprising 320 proteins for diverse eukaryotic lineages (733) was used as starting point for data set construction^7^. With a few exceptions that were kept as outgroup, non-fungal linages were removed. The data set was then expanded by adding a broad range of fungal lineages as follows: 1) Protein sequences for each taxon were clustered with CD-HIT v. 4.8.1^45^ using an identity threshold of 85%. 2) Candidates for homologous copies were retrieved by BLASTP searches^43^ using the 320 proteins as queries (e-value: 1e-20; coverage cutoff: 0.5). 3) In three rounds, phylogenetic trees were inferred and carefully inspected in order to detect and remove paralogs and contaminants and select orthologs. In the first round, sequences were aligned with MAFFT v. 7.475^48^ using the --auto option, filtered with trimAL v. 1.4^55^ using a gap threshold of 0.8, and ML trees were inferred with FastTreeMP v. 2.1.11^46^ using -lg -gamma and options for more accurate performances. In the second and third round, non-homologues sequence stretches were filtered out prior to aligning using Prequal v. 1.02^26^ with a threshold of 0.95. Protein sets were then aligned using MAFFT G-INS-i with the following options: --allowshift, --unalignlevel 0.6, --maxiterate 0. Further non-homologues residues were removed with Divvier v. 1.01^27^ employing the -partial flag and alignments were trimmed with trimAL using a soft gap threshold of 0.05. Finally, partial sequences belonging to the same taxon that did not show evidence for paralogy or contamination were merged. Single-protein ML phylogenies (Data S1) were reconstructed with IQ-Tree v. 1.6.12^47^ under BIC-selected models including site-homogeneous models (such as LG) and empirical profile models (C10–C60) and node support was inferred by ultrafast bootstrap approximation^59^ (1,000 replicates).

Of the initial 320 proteins, 21 were removed due to a high proportion of missing taxa or incongruous clustering of several clades. The final data set comprised 299 proteins and 637 fungal taxa plus 24 non-fungal taxa (Data S1). The protein sequences were concatenated using SCaFoS v. 1.25^51^. The obtained matrix (129,814 amino acid positions) was then used to calculate an initial ML tree with IQ-Tree employing the site-homogeneous model LG+G+F and ultrafast bootstrap approximation (1,000 replicates; Figure S1). To allow computationally more demanding analyses, a taxon-reduced data set was built representing all major fungal groups and discarding taxa with more missing data. Corresponding sequences were newly aligned, filtered, and concatenated (same settings as described above; 101 taxa, 128,450 amino acid positions). New ML trees were inferred for each of the taxon-reduced protein alignments as described above (Data S1).

#### Phylogenomic analyses

The taxon-reduced matrix was used to reconstruct a ML tree with IQ-Tree using the best-fitting site-heterogeneous model LG+C60+G+F with the PMSF approach^16^ to obtain nonparametric bootstrap support (100 replicates; Figures 1B and S3). Copies of this tree were manually edited to test alternative topologies with the approximately unbiased test (AU test^60^). The taxon-reduced matrix was also subjected to Bayesian analyses using PhyloBayes-MPI v. 1.8^50^ with the site-heterogeneous CAT+GTR+Γ4 model and the -dc option to remove constant sites. Three independent Markov Chain Monte Carlo (MCMC) chains were run for 1,600 generations (all sampled). For each chain, the burnin period was estimated by monitoring evolution of the log-likelihood (Lnl) at each sampled point, and 750 generations were removed from all chains. A consensus tree (Figure 1A) was built and global convergence between chains was assessed with PhyloBayes’ built-in tools. As frequently recognised in PhyloBayes, global convergence was not received in all combinations of the chains with maxdiff = 1 and meandiff = 0.0167504. To discriminate whether potential discrepancies between this tree and the tree calculated by ML are due to the usage of different software packages or due to different evolutionary models, a further Bayesian tree (Figure S5) was inferred under the LG+C60 model (-dc -catfix C60 -lg -dgam 4; three chains, each with 2,000 generations; convergence was not received; maxdiff = 1, meandiff = 0.0268007). Posterior predictive analyses were then run for both tree inferences in order to informatively select the best topology.

The single-protein ML trees of both the full and the taxon-reduced data sets (Data S1) were also analysed under the multi-species Coalescent (MSC) model in ASTRAL-III v. 5.7.7^42^ to incorporate protein tree uncertainties (Figures 1, S2, and S4). Prior to that, branches with bootstrap support <10 were collapsed as recommended by the developers. Quadripartition supports were calculated from quartet frequencies among the set of input trees^61^.

#### Sites removal analyses

25% and 50% of the most heterogeneous sites were removed from the taxon-reduced matrix using the script alignment_pruner.pl (https://github.com/novigit/davinciCode/blob/master/perl). Bayesian trees (Figure 2) were inferred form the pruned alignments with the CAT+GTR+Γ4 model: 25% pruned: three chains, each with 1,600 generations (burnin 1,000); convergence was not received; maxdiff = 1, meandiff = 0.0187389. 50% pruned: three chains, each with 2,450 generations (burnin 1,250); convergence was not received; maxdiff = 1, meandiff = 0.0128211. Also for the taxon-reduced matrix, fast-evolving sites were estimated with IQ-Tree under the LG+C60+G+F model, employing the -wsr flag and giving the PMSF tree as fixed tree topology. Sites were then removed in 10,000 increments and ML trees were inferred from the site-reduced alignments using IQ-Tree (LG+C60+G+F -bb 1,000 -bnni -wbtl; Figure 3).

#### Long branch attraction test

From the taxon-reduced data set, the eleven fastest-evolving lineages (two Sanchytriomycota species and nine Microsporidia species) were removed and the corresponding sequences were newly aligned, filtered, and concatenated as described above (90 taxa, 129,049 amino acid positions). The matrix was then used to infer a ML tree with IQ-Tree using the best-fitting LG+C60+G+F model and the PMSF approach to obtain nonparametric bootstrap support (100 replicates; Figure S6). Also, one further Bayesian tree was reconstructed using the CAT+GTR+Γ4 model (Figure S6): three chains, each with 900 generations (burnin 500); convergence was not received; maxdiff = 1, meandiff = 0.0188019.

### Quantification and statistical analysis

Phylogenetic model selection was based on IQ-Tree’s Bayesian information criterion^47^. Statistical support for phylogenies was obtained using non-parametric bootstraps, ultrafast bootstraps^59^, Bayesian posterior probabilities in BI^50^, and quadripartition support values in ASTRAL^42,61^. Alternative topologies were tested using the approximately unbiased test implemented in IQ-Tree^47^, and the fit of evolutionary models (*CATGTR* and *BI-C60*) was tested using posterior predictive analyses implemented in the PhyloBayes software package^50^.

