## Supplementary material for "Phylogenomics unravels the early diversification of fungi": Document_S1_Figures_S1-S6

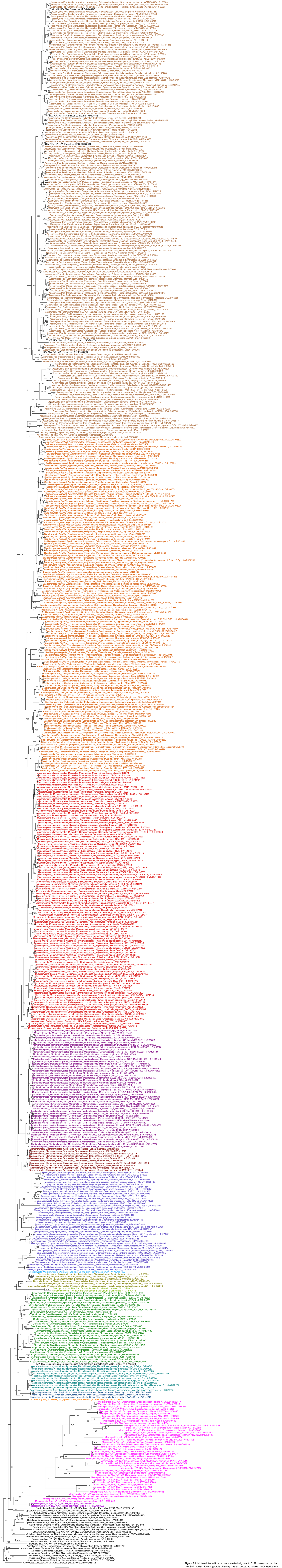

**Figure S1:** ML tree inferred from a concatenated alignment of 299 proteins under the LG+G+F model. Node support is given by ultrafast bootstrap values (1,000 replicates).

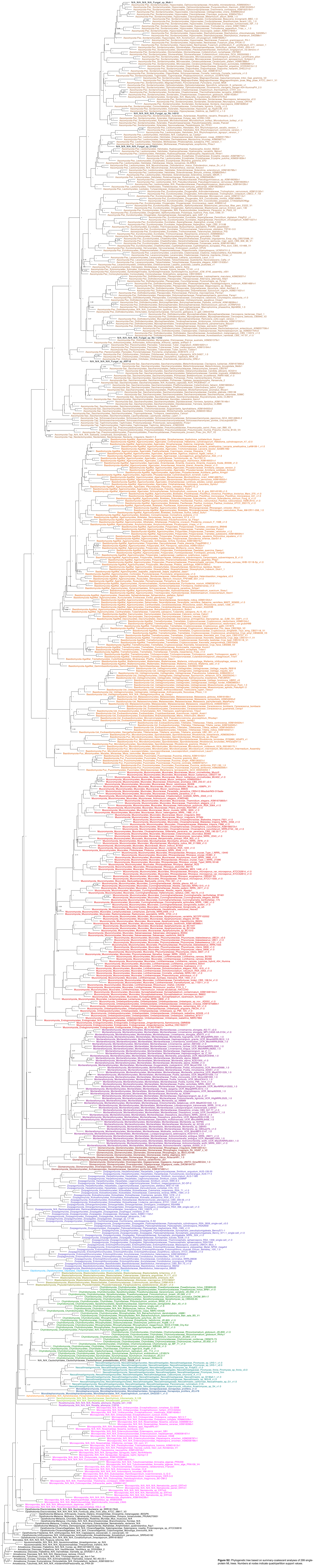

**Figure S2:** Phylogenetic tree based on summary-coalescent analyses of 299 single-protein ML trees. Numbers at nodes indicate quadripartition support values.

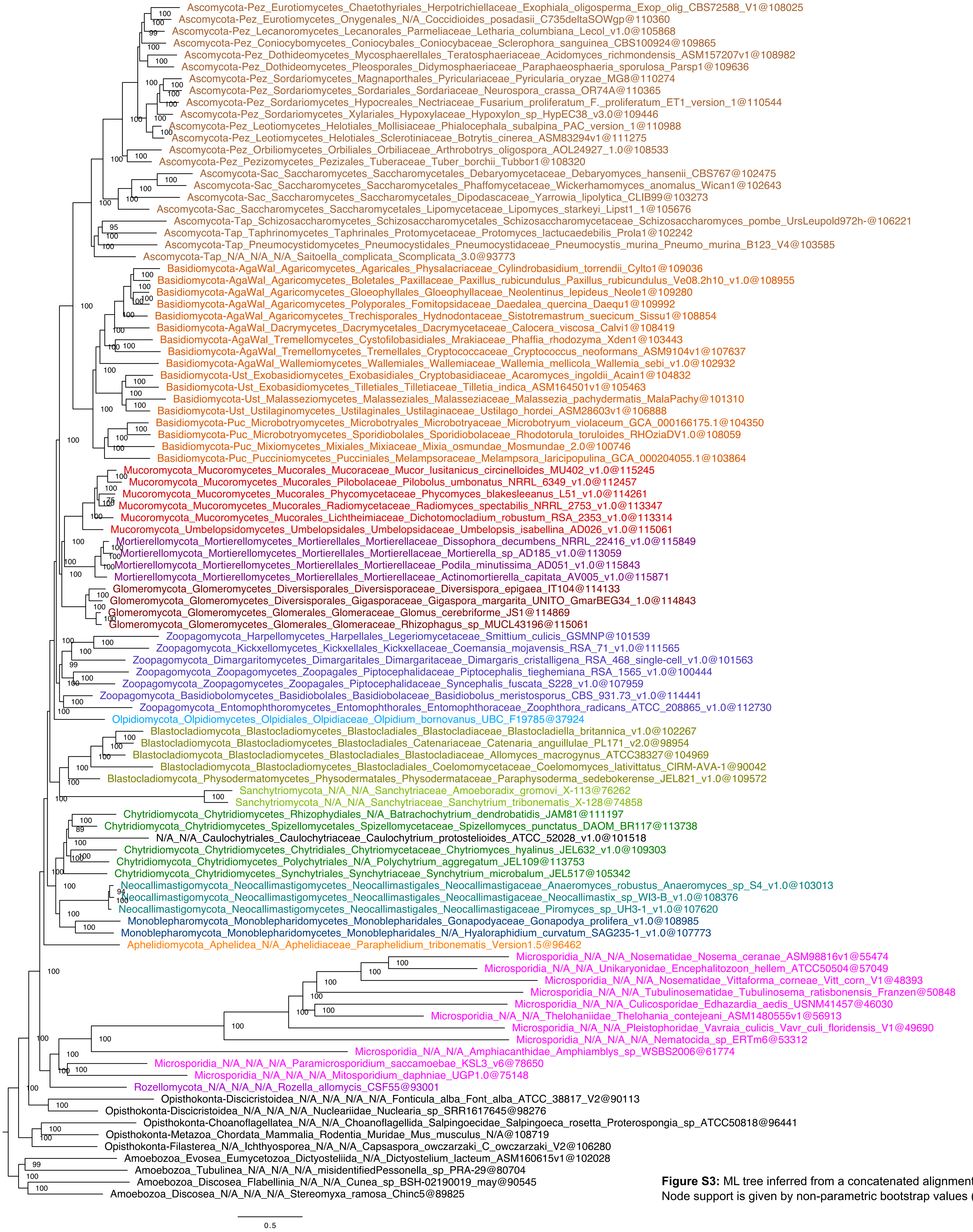


**Figure S3:** ML tree inferred from a concatenated alignment of 299 proteins under the LG+C60+G+F-PMSF model. Node support is given by non-parametric bootstrap values (100 replicates).

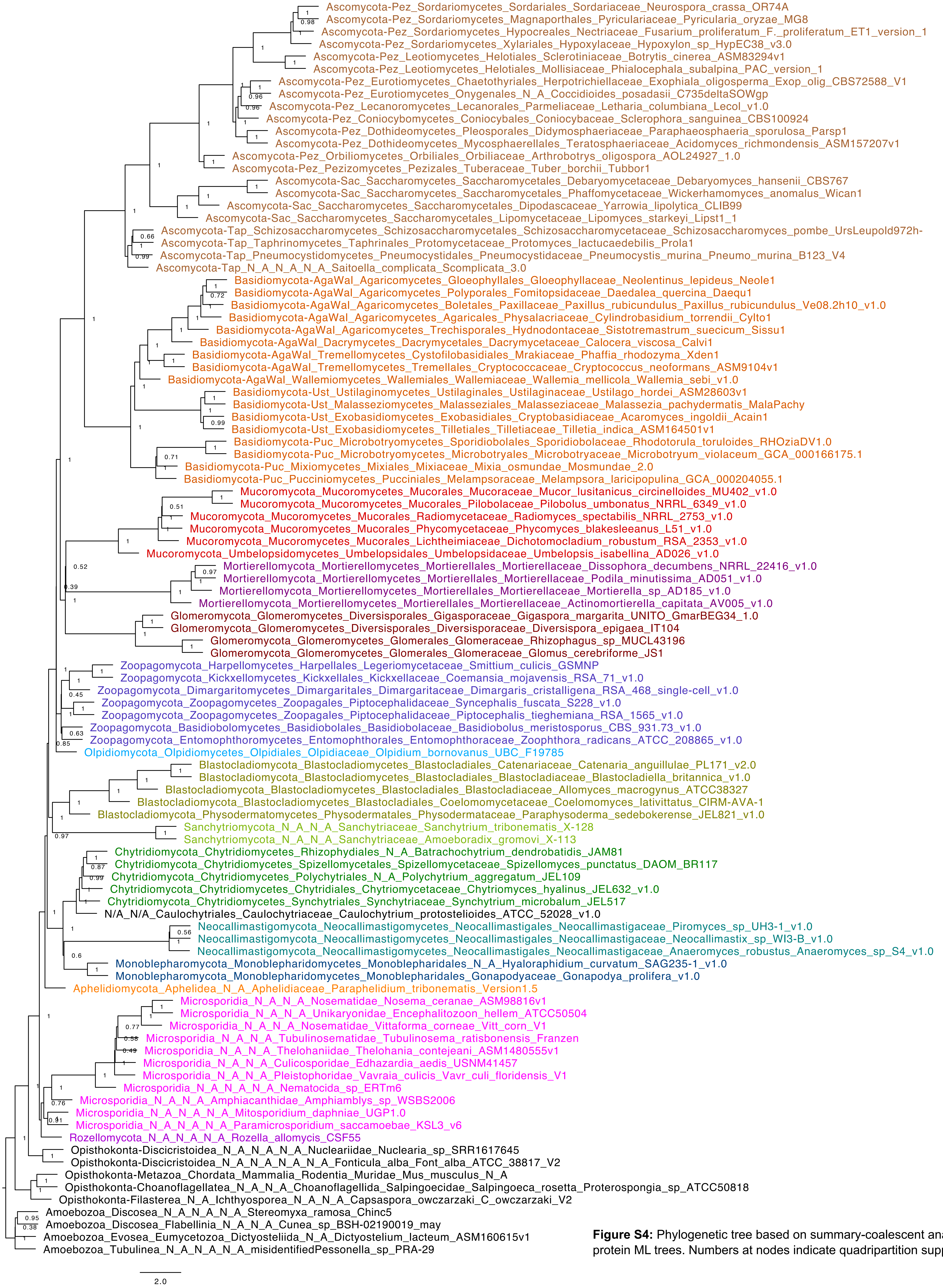


**Figure S4:** Phylogenetic tree based on summary-coalescent analyses of 299 single-protein ML trees. Numbers at nodes indicate quadripartition support values.

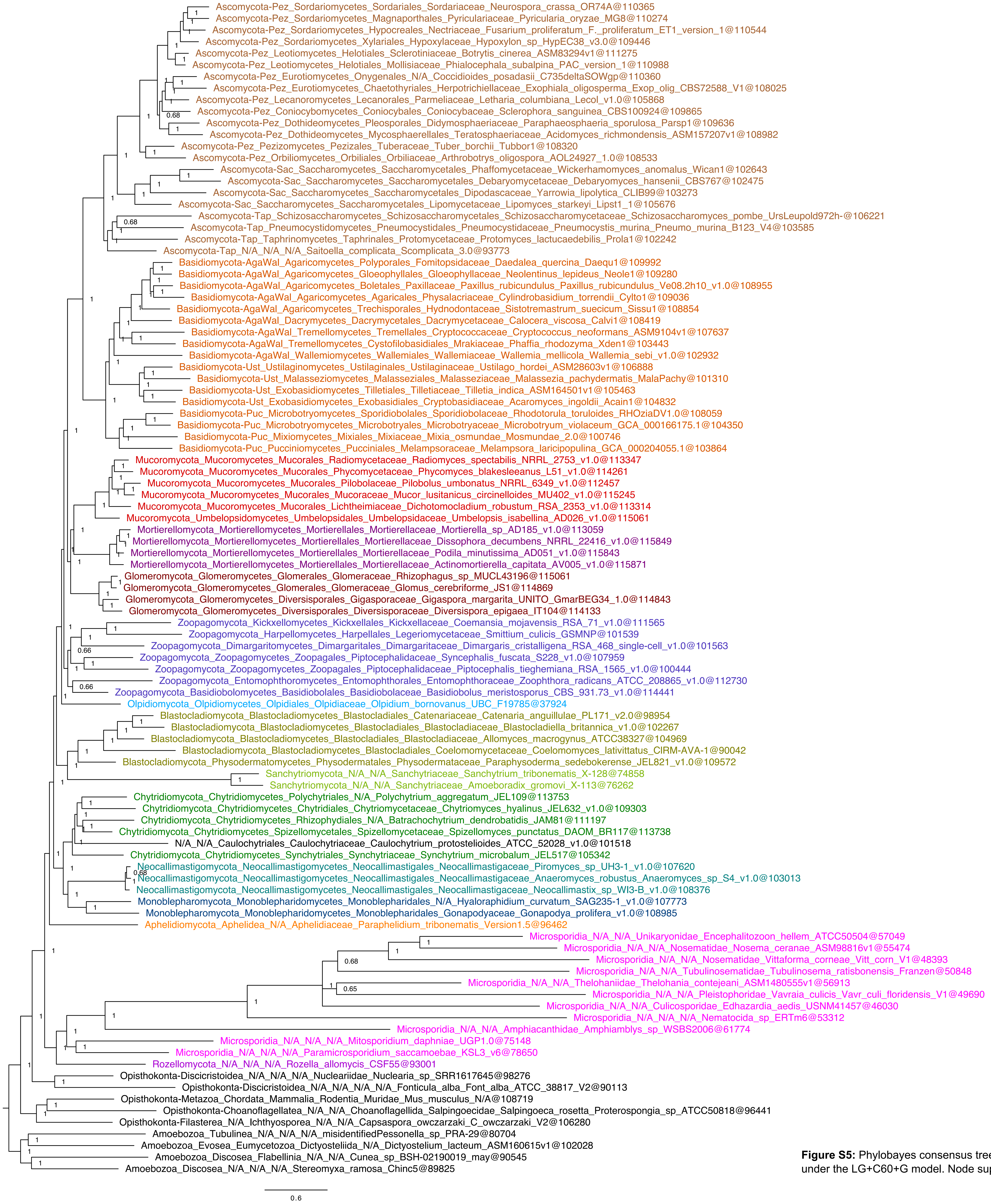


**Figure S5:** Phylobayes consensus tree inferred from a concatenated alignment of 299 proteins under the LG+C60+G model. Node support is given by Bayesian posterior probabilities.

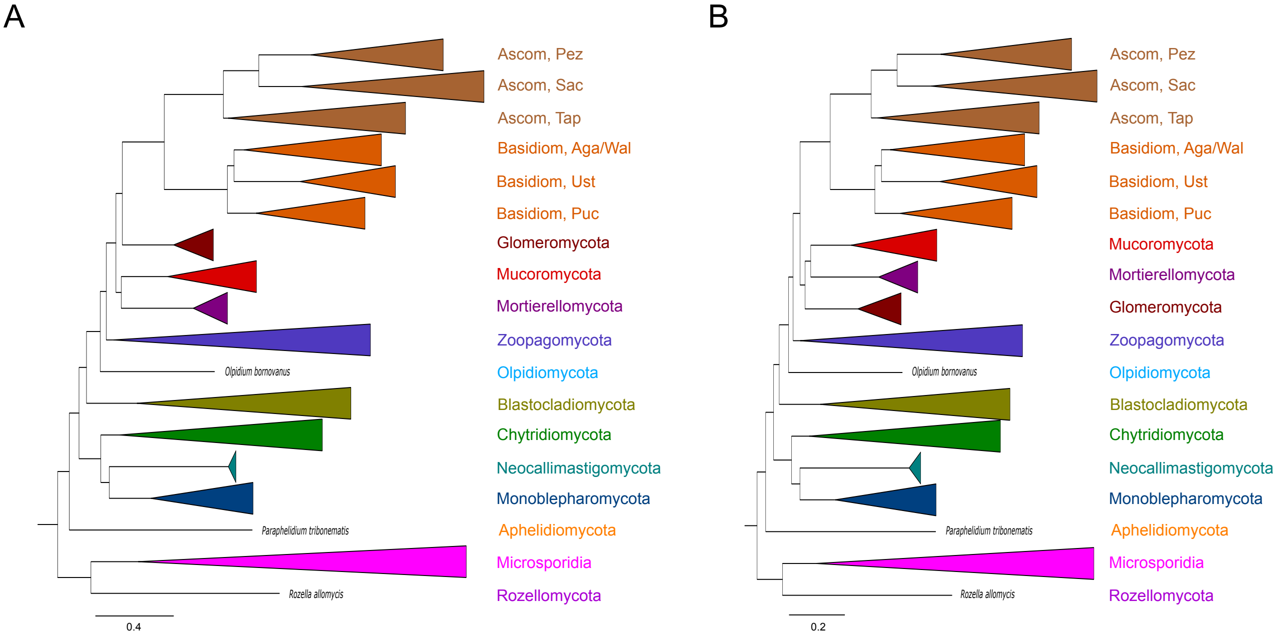


**Figure S6:** Phylogenetic trees inferred from a concatenated alignment (299 proteins) excluding the following fast-evolving species: Sanchytriomycota: *Amoeboradix gromovi* and *Sanchytrium tribonematis*; Microsporidia: *Nosema ceranae*, *Encephalitozoon hellem*, *Vittaforma corneae*, *Tubulinosema ratisbonensis*, *Edhazardia aedis*, *Thelohania contejeani*, *Vavraia culicis*, *Nematocida* sp., and *Amphiamblys* sp.
(A) Phylobayes consensus tree inferred under the CAT+GTR+Γ4 model. (B) ML tree inferred under the LG+C60+G+F-PMSF model. Support was maximal for all nodes shown in (A) and (B).
